# Development of a stable microbial community for the study of plasmid-mediated antimicrobial resistance in a continuous culturing device

**DOI:** 10.64898/2026.10.13.758845

**Authors:** Thu-Hien Vu, Tania Alonso-Vásquez, Elisa Sestini, Alessio Masoni, Katharina Grabowski, Camille Andrieu, Leanid Laganenka, Francesca Di Patti, Marco Fondi, Marco Galardini

## Abstract

Horizontal gene transfer (HGT) is a major contributor to the spread of antimicrobial resistance (AMR) in microbial communities, particularly through conjugative plasmids. The dynamics of the spread of these plasmids in a susceptible microbial community requires a defined and controllable *in vitro* system, which is currently lacking from existing laboratory evolution systems. Here we describe the development of TriComm, an engineered microbial community of three strains with complementary amino acid auxotrophies, maintained in continuous culture under real-time optical and fluorescence monitoring. We show that two conjugative plasmids carrying resistance genes (*bla*_OXA-48_ and *mcr-1*) generally rescued TriComm from collapse under antimicrobial treatment, and that this protection was dependent on conjugative transfer. Community composition remained stable across most conditions, including after the AMR plasmid had spread through the population. Through mathematical modeling we predict that modulating both drug efficacy and conjugation rate can lead to the collapse of the community, suggesting potential strategies to limit the spread of AMR via HGT.

## Introduction

Antimicrobial resistance (AMR) is a pressing public health problem causing a substantial infection burden worldwide^1^. AMR is driven not only by spontaneous mutations emerging in the bacterial genome, but also by the transfer of resistance-conferring genetic elements between clades. Compared to mutation, horizontal gene transfer (HGT) allows a bacterial host to acquire complete genes, thus potentially leading to a large shift in susceptibility to an antimicrobial. HGT is mainly driven by mobile genetic elements which can spread within and across bacterial species, allowing microbial communities to adapt faster than would be possible through mutation alone^2^. A quantitative and mechanistic understanding of the evolution of AMR therefore requires experimental systems that consider the horizontal transmission of resistance genes.

Conjugative plasmids are a special class of mobile genetic elements: they encode a complex molecular machinery that recognizes plasmid-free cells and mediates its own transfer into them^3^. The dynamics of plasmid spread are complex, shaped by the fitness cost of carrying the plasmid, its copy number, its conjugation rate, and the selective advantage conferred by the AMR genes it encodes^4^. In the context of antimicrobial treatment these factors, together with treatment strength and dynamics, determine whether a susceptible microbial community can be rescued by the presence of donor cells carrying an AMR conjugative plasmid^5,6^. Combining empirical measurements with mathematical modeling can therefore help elucidate the dynamics of AMR spread via conjugation.

Two clinically relevant classes of conjugative resistance plasmids illustrate this complexity. pOXA48 plasmids encode the *bla_OXA-48_* carbapenemase gene and can spread across a range of enterobacterial species^7,8^. This gene catalyzes the hydrolysis of β-lactam antibiotics; expressing cells can therefore provide a “public good” by reducing the effective antibiotic concentration, and thus potentially allowing susceptible cells to survive treatment^9–11^. This can reduce the selective pressure that favors plasmid spread, especially when the fitness cost of carrying the plasmid is high relative to the protection it confers on the wider community. Plasmids carrying the *mcr-1* gene, such as IncX4 plasmids, instead confer resistance to polymyxins such as colistin by modifying the cell envelope, conferring no benefit to the rest of the community^12,13^. These two plasmid types therefore might give rise to distinct spread dynamics under antimicrobial treatment, despite posing a similarly serious threat to public health. Comparing them offers an opportunity to derive generalizable principles governing AMR spread via HGT.

Despite the relevance of HGT and the need to develop a quantitative understanding of this process, *in vitro* laboratory evolution has historically been skewed toward the study of clonal populations in which adaptation can only emerge via mutation. More recently, a number of experimental studies have shown that HGT can be studied in a controlled environment, demonstrating its role in adaptation to selective pressures and deriving quantitative models of its impact on microbial communities^14–16^. Despite these advances, the impact of conjugative plasmid spread on microbial community dynamics has yet to be studied within an *in vitro* laboratory evolution setting that would allow for the development of a quantitative model. Existing assays such as conjugation rate measurements by selective plating are useful, but do not capture community dynamics and the balance between selective advantage and fitness cost, which can be revealed by a growth-based assay. Serial passaging is the *de facto* standard approach in laboratory evolution, but the imposition of population bottlenecks at each passage risks affecting community composition and dynamics^17^. Continuous liquid culturing would therefore be an ideal setting to study the dynamic of a microbial community under antimicrobial treatment and in the presence of an AMR conjugative plasmid that might rescue it. Better still, a continuous culturing system capable of real-time growth monitoring would provide the quantitative data needed to model these dynamics.

The other element required to build an experimental system in which to study HGT dynamics is a defined and stable microbial community. In the absence of interactions between the community members, the faster growing member would outcompete and exclude the others. Metabolic interdependencies, a form of positive interactions, have been shown to produce robust and stable microbial communities^18^. Furthermore, auxotrophic interactions, in which one member depends on a metabolite produced by another member, can be designed to be intertwined, so that the removal of any single community member leads to the collapse of the whole community^19–21^. This in turn provides an ideal system in which to test the spread of an AMR plasmid under antibiotic treatment.

Here we describe the development and testing of TriComm, a community consisting of three engineered *E. coli* BW25113 strains whose complementary amino acid auxotrophies make, in theory, each member essential for community survival in minimal media^19^. We monitored TriComm growth in real time using the Chi.Bio culturing system^22^, which allowed us to track community composition through fluorescence measurements. We validated this real-time tracking approach against colony counting and flow cytometry. In the absence of antimicrobial treatment or conjugative plasmids, we observed the long-term persistence of all three TriComm members with periodic oscillations. We then introduced a pOXA48 conjugative plasmid carrying the *bla_OXA-48_* carbapenemase gene^23^ and an *mcr-1*-carrying conjugative plasmid (PN23^24^) into one member of the community, and recorded community dynamics in the presence or absence of the cognate antibiotic. We used the resulting data to build a mathematical model to investigate the tension between plasmid-mediated rescue and community collapse.

## Results

### Development of TriComm, a synthetic obligate syntrophic microbial community

Our first objective was to develop a microbial community in which individual members depended on each other for growth; in this scenario, a selective pressure that removes one or more members would lead to the community’s collapse. In such a community, the introduction of a horizontally transferable resistance determinant would promote community survival, although with an unknown impact on community dynamics and composition.

We constructed a synthetic community of three members, each carrying two gene knockouts in amino acid biosynthetic genes, arranged such that each member would depend on the other two to obtain the amino acids it could not synthesize itself. We named this three-strain community TriComm (**Figure 1A**). We chose three amino acid biosynthetic genes (*metA*, *lysA*, and *pheA*) and generated three double knockout strains in an *E. coli* BW25113 background: PM (Δ*pheA*, Δ*metA*), PL (Δ*pheA*, Δ*lysA*), and LM (Δ*lysA*, Δ*metA*). This choice was based on a previous study which identified this combination as resulting in a high growth yield in minimal media lacking the corresponding amino acids^19^. As the constructs from the original study were not available, we generated new ones from single gene knockout strains in the KEIO collection^25^, and confirmed them by whole genome sequencing (**Supplementary Figure 1**).

**Figure 1.**
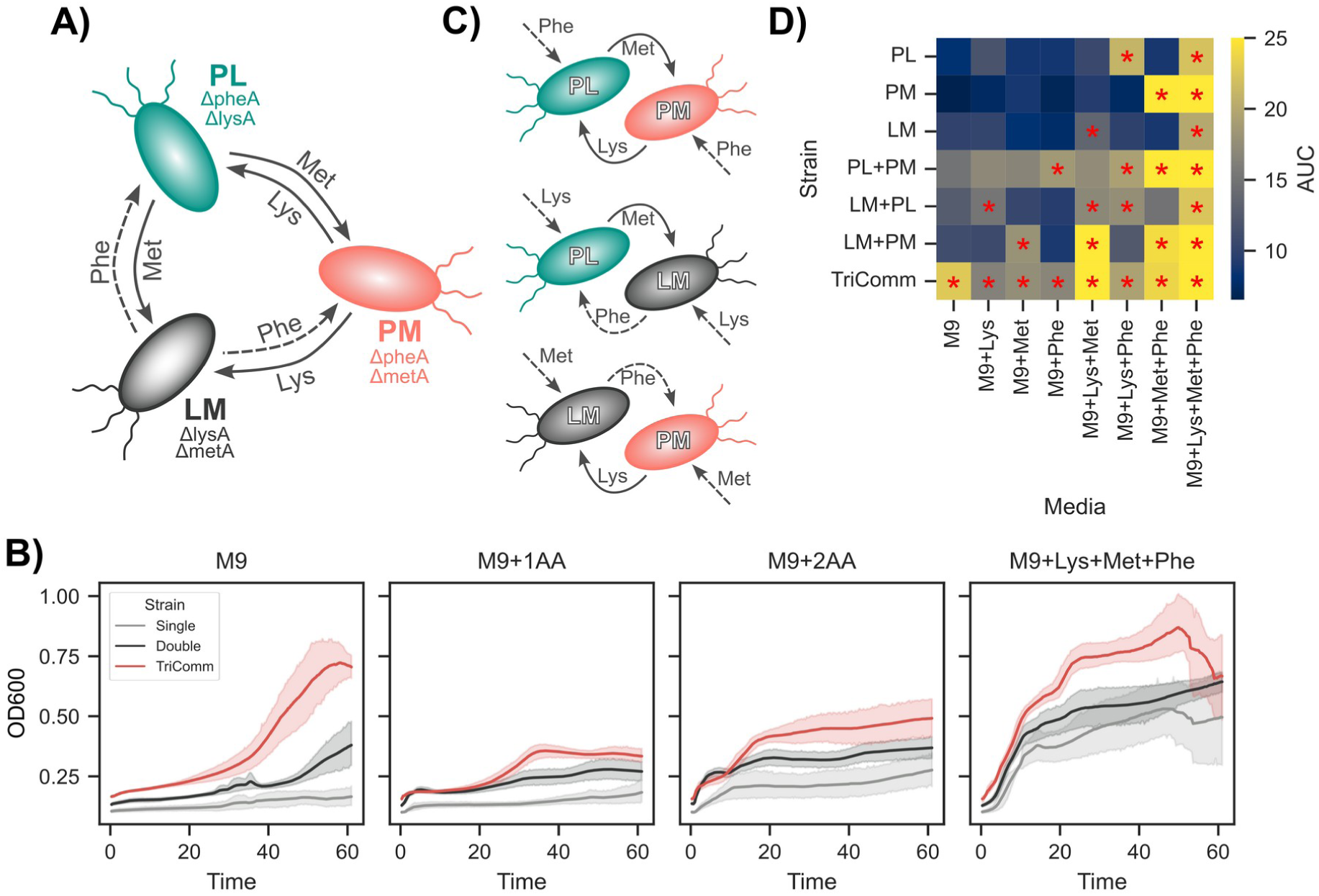
Characterization of TriComm, a stable synthetic microbial community. A) Schematic representation of TriComm’s members gene knockout profiles and interdependencies of TriComm members. The dashed line indicates the theoretical phenylalanine auxotrophy. B) Growth curves of the single strains, pairwise combinations, and the full TriComm community across different minimal media compositions; without added amino acids (M9), with one of the three key amino acids added (M9+1AA), with two amino acids added (M9+2AA), and with all three amino acids added (M9+Lys+Met+Phe). The solid line indicates the mean across replicates and the shaded line the 95% confidence interval. Time is expressed in hours. C) Schematic of interdependencies between pairs of strains, indicating which amino acid is theoretically required for growth for each two-member community (dashed arrows). D) Mean area under the curve (AUC) across replicates for each community grown in each media composition. Red asterisks indicate the community and media pairs in which we expected to observe significant growth. Lys: Lysine, Met: methionine, Phe: phenylalanine.

We then tested whether the individual strains (*i.e.* LM, PL, and PM) and two member communities (*i.e.* PL+PM, LM+PL, LM+PM) were indeed unable to grow in minimal media lacking the amino acids lysine, methionine, and phenylalanine. As expected, neither the single strains nor the two-member communities grew significantly in the absence of their required amino acids, while TriComm reached a significantly higher OD, comparable to that observed for TriComm grown in media containing all three amino acids (**Figure 1B**). Unexpectedly, TriComm grew to significantly lower yields and only marginally more than the other strains in minimal media supplemented with one or two amino acids. We observed that, especially in the case of minimal media with two amino acids, growth was characterized by an early but stunted exponential phase, likely reflecting transitory growth of one or two members at the expense of the rest of the community (**Figure 1B** and **Supplementary Figure 1**). For the single strains and two-members communities, we observed growth yields comparable to those of TriComm only when all three amino acids were provided.

Having validated that TriComm was the only community able to grow in minimal media without amino acids, we next sought to test the robustness of the auxotrophies generated via gene knockouts by growing the three individual strains and the three possible 2-members communities. The individual strains were predicted to grow only in the presence of the two amino acids corresponding to the two knocked out biosynthetic genes, while for each of these two-members communities were predicted to depend on a single amino acid for growth: phenylalanine for PL+PM, lysine for LM+PL, and methionine for LM+PM (**Figure 1C**). As expected, we observed appreciable growth (area under the growth curve > 12.5) of the individual strains only in the presence of their corresponding amino acids: lysine and phenylalanine for PL, methionine and phenylalanine for PM, and lysine and methionine for LM (**Figure 1D**, red asterisks). Similarly, we observed growth for all three two-members community in media only containing their missing amino acid, and lower AUC values otherwise; the exception was the PL+PM community, for which we observed an AUC > 15 in any media that contained at least one amino acid, rather than exclusively in those containing phenylalanine (**Figure 1D** and **Supplementary Figure 1**). This unexpected result suggests that the phenylalanine auxotrophy may be more “leaky” than the other two, potentially resulting in a lower relative proportion of the phenylalanine producing strain (LM) during sustained TriComm growth. We further tested this hypothesis in continuous culture, and we observed that indeed the PL+PM community and TriComm grew with a similar dynamic in minimal media (**Supplementary Figure 2**). TriComm is therefore a synthetic community in which two of its members are essential (*i.e.* PL and PM), and one (*i.e.* LM) is not and depends on the other two.

### Tracking of community composition in a continuous culturing device

We engineered TriComm to also enable tracking of changes in its composition over time. We chose to culture TriComm in the Chi.Bio device^22^ for two reasons: first, because it allows continuous culturing, thus removing population bottlenecks and excessive manual labor, and second, because the Chi.Bio bioreactors are equipped with a variety of sensors, including an optical density sensor and a fluorescence sensor, each taking a measurement every minute. We therefore introduced two small non-mobilizable plasmids expressing a fluorescent marker in two of the three TriComm members: one expressing GFP in PL, and one expressing mScarlet in PM (**Figure 1A**).

We then tested the ability of the Chi.Bio device to track the composition of communities with a predefined ratio of each member, as well as with two other methods (plate counting and flow cytometry), each requiring extracting a sample from the bioreactor (**Figure 2A**). We chose a negative control (M9 minimal media), individual members in isolation (PL, PM, and LM), TriComm communities with two minority members at 1% of the inoculum size of the majority member (PLpmlm, plPMlm, and plpmLM), TriComms with two majority members (PLPMlm, PLpmLM, and plPMLM), and a balanced TriComm (PLPMLM). The Chi.Bio fluorescent sensor could identify the corresponding fluorescent marker in single member and two minority members communities, whereas the sensitivity was somewhat reduced for two majority members and balanced communities (**Figure 2B**, left panel), likely due to the dampening or scattering of the fluorescent signal by the cells of the other community members. We performed colony counting identifying the individual members by the color of the colony on kanamycin selective plates, and we observed a generally accurate, although noisy, estimation of the minority isolates (**Figure 2C**, left panel). Strain LM was underestimated in both communities with one majority strain (PLpmlm and plPMlm), and could not be identified by counting in the PLPMlm community; we hypothesized that the smaller colony size of strain LM (**Supplementary Figure 3**) contributed to this underestimation. When we measured the same communities using flow cytometry, we observed the opposite phenomenon, namely a slight overestimation of the proportion of the LM strain in all communities in which it was present as the minority strain ( **Figure 2D**, left panel). The most likely explanation for this phenomenon is the presence of cells belonging to the PL and PM strains not expressing the fluorescent marker, which we estimate to comprise about 2% of total cells on average. In order to compare the measurements obtained by the Chi.Bio fluorescent sensor with those of the other two methods, we used the ratio of the green over the red signal (**Figure 2B-D**, right panels). We observed that for both the Chi.Bio and flow cytometry measurements, the ratios obtained from single member green (*i.e.*, PL) and red (*i.e.*, PM) communities provided a limit of detection for these two methods, since the other strain was absent in those communities; plate counting was not subject to this limitation, as we did not identify any colonies of the opposite color on the single member samples. All three methods yielded a consistent green-to-red ratio in communities where PL and PM were present at the same ratio (*i.e.* plpmLM, PLPMlm, and PLPMLM), as well as those in which one member was the minority with respect to the other strain. From these experiments we concluded that either of these three methods could identify shifts in the relative proportion of the PL and PM strains with a lower bound of 1%, and that plate counting underestimates the LM strain, while flow cytometry overestimates it.

**Figure 2.**
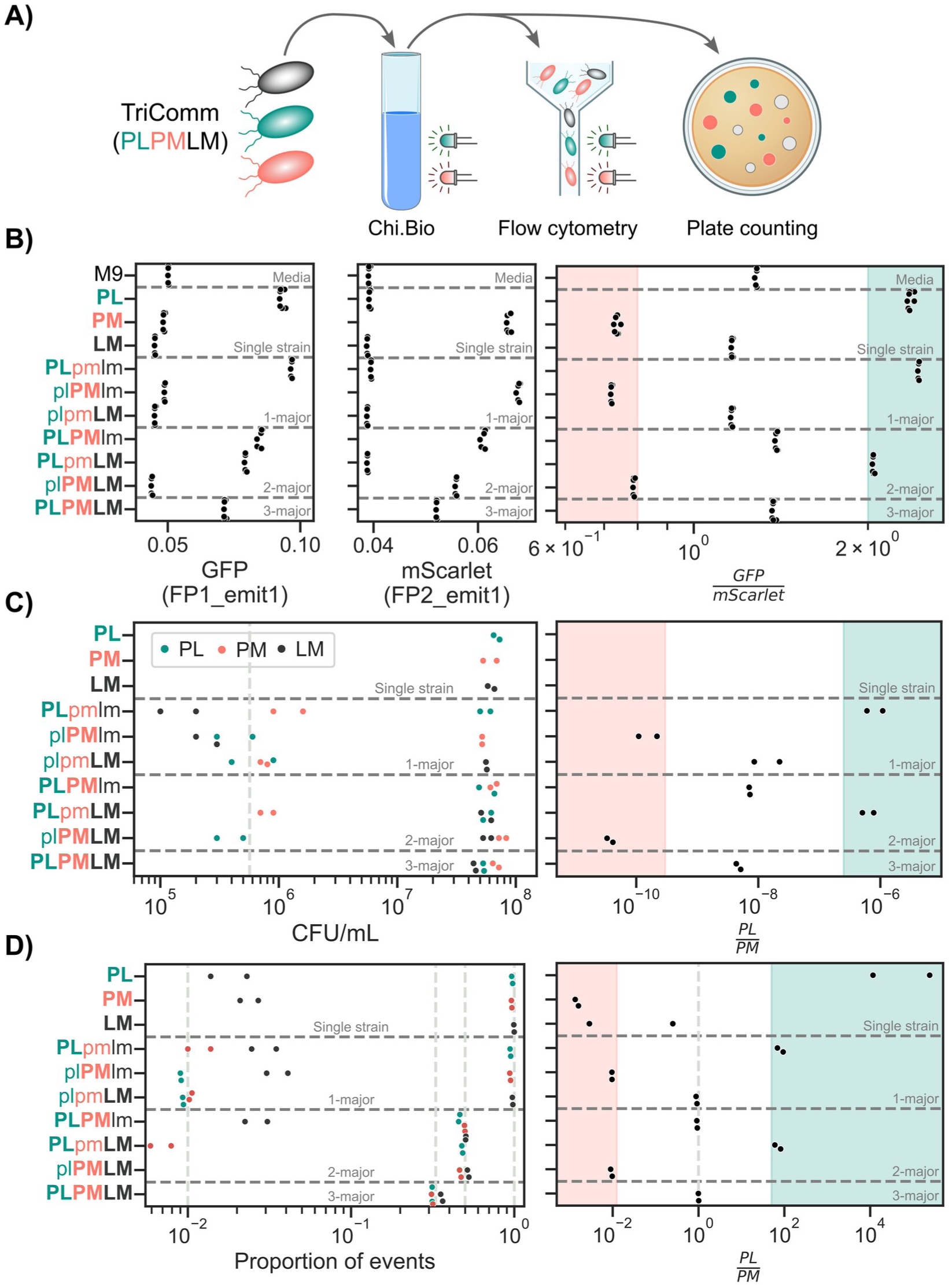
Sensitivity of three strain detection methods on defined TriComm communities. A) Schematic representation of the sensitivity experiments and of the three detection methods. B) Fluorescent signals recorded by the Chi.Bio device across two wavelengths (Methods). C) Measured CFU/mL from colony counting from plates inoculated with samples from each community. D) Proportion of events belonging to each category (GFP, mScarlet, and blank) from samples analyzed through flow cytometry. Left side of panels B to D show the raw data, while the right panels show the green to red signal ratios.

We then tested our ability to track the composition of a TriComm community grown continuously in a Chi.Bio bioreactor, using all three methods across three biological replicates (**Figure 3**). We observed a reproducible oscillatory behavior in the overall community, as measured by optical density and plate counting (**Figure 3A-B**), which persisted up to 1000 hours of continuous growth (**Supplementary Figure 4**). These oscillations were also evident in the Chi.Bio fluorescent measurements, potentially indicating variations in the relative proportions of the PL and PM strains. We indeed observed that the relative proportions of the three isolates changed over the course of continuous growth (**Figure 3B-C**); both plate counting and flow cytometry showed a sharp reduction in the levels of the LM strain, from roughly a third of the community to below 1%, which we had previously determined to be the limit of detection for both methods. This confirms that strain LM, while dependent on the other two isolates, is dispensable for the overall community. Strains PL and PM also showed changes in their relative ratio using the three methods, although not as dramatically as the LM strain and generally within the bounds we had defined in our earlier mixing experiments (**Figure 3B-F**). Taken together, these experiments suggest that TriComm is a stable community in which each member oscillates around a reproducible level, at least in the relatively long time frame used in these culturing experiments.

**Figure 3.**
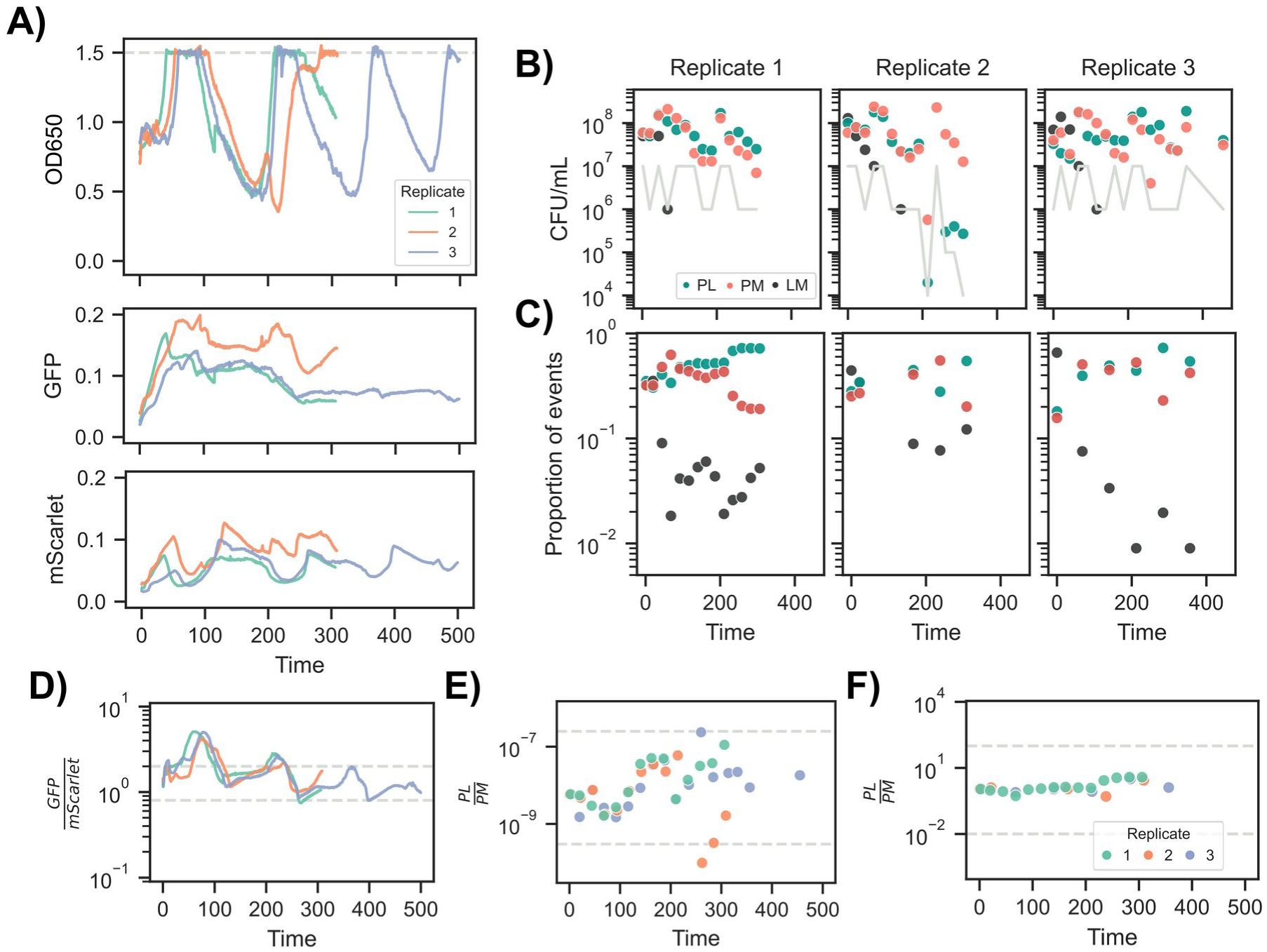
TriComm community dynamics during continuous growth. A) Optical density and fluorescence measurements taken at one minute intervals by the Chi.Bio device. The horizontal dashed line indicates the OD level at which the bioreactor switches from chemostat to turbidostat. B) Colony forming units for the three strains from daily samples; the grey line indicates the limit of detection for the minority strain. C) Proportion of events belonging to each strain for samples analyzed through flow cytometry. D-F) Ratio of green and red signals from the bioreactor (D), plate counting (E), and flow cytometry (F). Dashed grey horizontal lines indicate the values at which PL (top line) and PM (bottom line) are predicted to be at 1% of the abundance of the other strain.

### Studying dynamics of plasmid-mediated spread of antimicrobial resistance

Having established that TriComm remains stable over a long time period, and that we could track its composition, we then set out to study how it would respond to an antimicrobial selective pressure, and how a conjugative plasmid conferring resistance would spread with or without such pressure. We decided to use two clinically relevant plasmids, pOXA48 and PN23, conferring resistance to penicillins/carbapenems and polymyxins, respectively. For pOXA48 specifically, we used two plasmid variants: a completely functional variant, PVI, and a non-conjugative variant lacking the conjugation machinery, PVB^23^. For all plasmids (*i.e.*, PVI, PVB, and PN23), we introduced them into the LM strain and grew the resulting community with or without the corresponding antibiotic (*i.e.*, ampicillin for PVI/PVB, and colistin for PN23, using two concentrations for the latter), comparing them against a community without any AMR plasmid (TriComm KAN, **Figure 4A**). We also used plate counting in the presence of the relevant antibiotic to identify TriComm transconjugants and how their abundance changed over time. We chose to use strain LM as the donor as the removal of the other two strains through antibiotic treatment would lead to its extinction, thus making conjugation essential.

**Figure 4.**
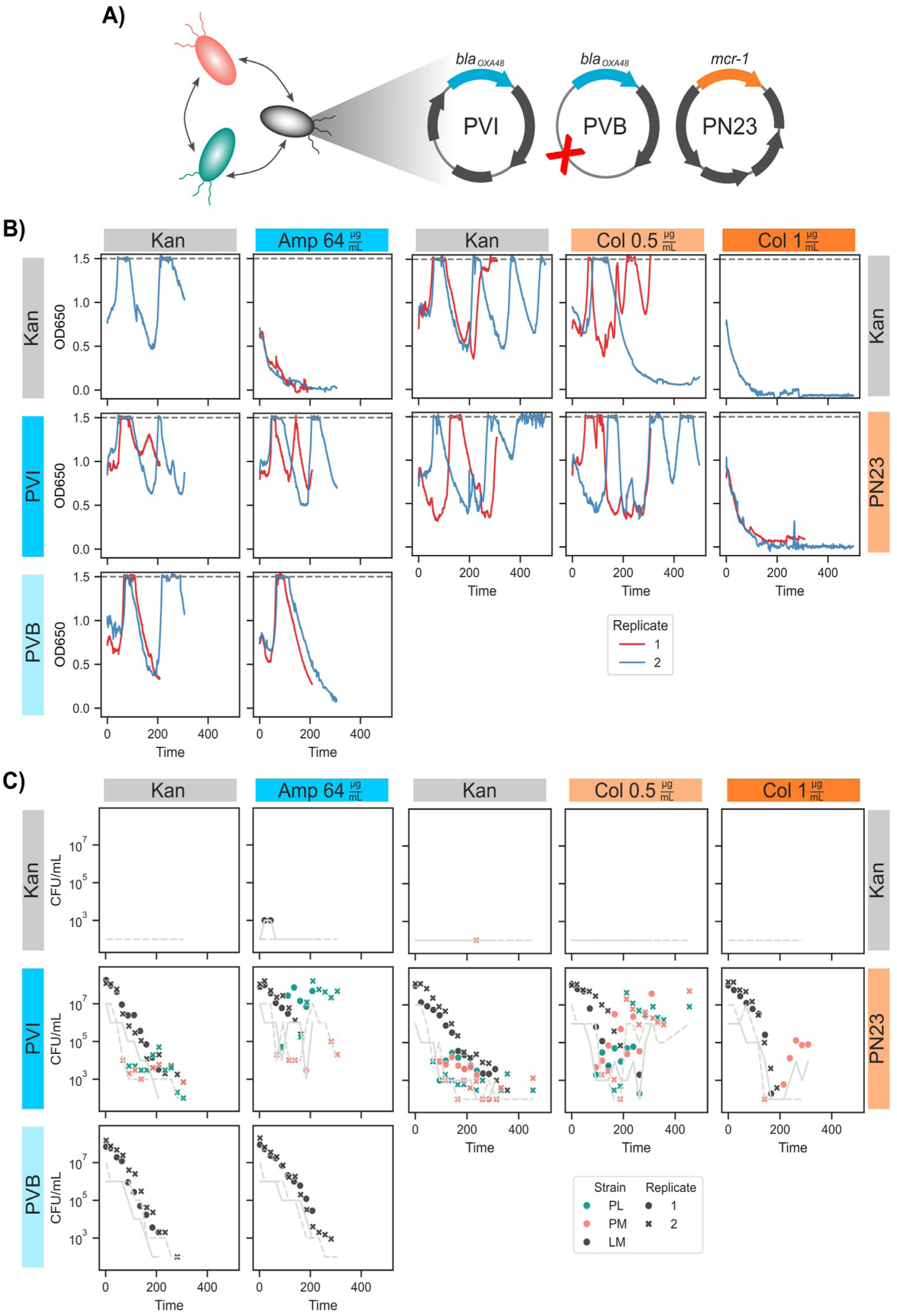
TriComm community dynamics under different selective pressures and conjugation regimes. A) Schematic of the different plasmids used in the growth experiments. The red cross on the PVB plasmid indicates the deletion that renders this plasmid unable to transfer via conjugation. B) Optical density measurements for each TriComm, selection and plasmid combinations, and across two biological replicates. The horizontal dashed line indicates the OD level at which the bioreactor switches from chemostat to turbidostat. C) Colony forming units for the three TriComm members on selective media containing either ampicillin (two leftmost columns) or colistin (three rightmost columns). The grey line indicates the limit of detection for the minority strain(s) at each time point.

Overall, we observed that, as expected, TriComm KAN’s growth was affected in the presence of both antibiotics; the decline in population numbers was particularly evident in the presence of ampicillin (**Figure 4B**). In the case of colistin, the impact of the drug on TriComm KAN was less pronounced across the two tested concentrations, with one replicate being able to grow with colistin at 0.5 µg/mL. We did not identify any colonies on colistin-selective media for this replicate, suggesting a transient non-genetic resistance mechanism or a lower than expected effective drug concentration (**Figure 4C**).

The presence of either AMR plasmid did not seem to affect growth in the absence of antibiotics, with all plasmid-carrying communities (TriComm PVI, PVB, and PN23) exhibiting oscillatory behavior similar to that of as TriComm KAN (**Figure 4B**). As expected, the presence of the AMR plasmid allowed the community to survive in the presence of the antibiotic, again showing a growth dynamic similar to TriComm KAN in regular media, with two exceptions. Growth in the presence of 1 µg/mL of colistin led to an apparent collapse of the TriComm PN23 community, suggesting that this drug concentration could not be counteracted by the activity of the *mcr-1* gene. The other exception was TriComm PVB grown in the presence of ampicillin, for which we observed community collapse after an initial peak in density (**Figure 4B**); we hypothesized that the inability of PVB to transfer to the other TriComm members could be responsible for this difference compared to TriComm PVI and PN23.

The optical density data collected by the Chi.Bio sensor showed that the presence of an AMR plasmid could keep the community stable when the selective pressure was not too high (*i.e.* as with TriComm PN23 at 1 µg/mL colistin), but not when the conjugative machinery was absent. This suggests that the AMR plasmid must spread to the essential strains ( *i.e.* PL and PM) for the TriComm community to survive, since their extinction would remove the production of essential amino acids from the rest of the community. We verified this hypothesis by plating samples taken over the course of the experiment on antibiotic-containing media; for TriComm PVI, PVB, and PN23 we expected to observe only LM colonies at the beginning of the experiment, regardless of antibiotic treatment. As expected, we found no LM colonies in TriComm KAN communities and a high LM cell density (> 10^7^ CFU/mL) for the other communities (**Figure 4C**). As hypothesized, we observed the appearance of green (*i.e.* PL) and red (*i.e.* PM) colonies on the selective plates after 68 to 184 hours of growth in all TriComm PVI and PN23 experimental runs except one, indicating successful transfer of the conjugative plasmid within the community, and which we verified by whole genome sequencing of 17 individual colonies isolated from the bioreactors (**Supplementary Table 1**). The only run in which we did not record a transconjugant was the first replicate of TriComm PVI grown without ampicillin. Read coverage across the chromosome and the four plasmid references confirmed the expected auxotrophy deletions and fluorescent markers in every sequenced clone (**Supplementary Table 1**). No known antimicrobial-resistance mutations were detected in any sequenced clone or in whole culture samples (**Supplementary Table 2**, and **Supplementary Discussion**). Despite conjugative transfer being observed consistently in PVI and PN23 communities, we recorded an overall log-linear reduction (−1.4 to −2.3 log₁_0_ CFU/mL per 100 hours) in colony counts for the communities grown in the absence of ampicillin or colistin, suggesting that the combination of the plasmid’s fitness cost and absence of selection drives the plasmid out of the community. Similarly, we recorded a log-linear decrease (−1.6 to −2.4 log₁_0_CFU/mL per 100 hours) in LM colonies on the selective plates under both growth conditions for TriComm PVB, which in the case of the ampicillin community confirms our hypothesis regarding the role of horizontal gene transfer in the survival of the community.

### TriComm composition remains largely stable

We next asked whether the effect of the antibiotics on community members and the transfer of the AMR plasmid led to significant changes in community composition. Although we anticipated that all three members should be present in the communities that did not decline, we also considered that larger shifts in the relative ratios of the TriComm members were possible. We used the high-frequency data collected by the Chi.Bio bioreactor (**Figure 5A-B**), and in particular the ratio between the GFP and mScarlet fluorescent signal ( **Figure 5B**) to identify likely shifts in community composition, which we could then validate using plate counting (**Figure 5C**) or flow cytometry (**Figure 5D**). We identified multiple instances in which the GFP to mScarlet ratio went beyond the threshold we had identified in our earlier community mixing experiments (**Figure 2**), indicating that either the PL or PM strain was reduced to a level below 1% of the other strain. In all but one growth condition ( *i.e.* TriComm PN23 with colistin at 1 µg/mL) we observed an early (∼100 hours) peak indicating that strain PL was present at a ratio of 1:0.01 with respect to strain PM; these observations were however generally not supported by the plate counting and flow cytometry data (**Supplementary Figure 5**), suggesting that these early peaks did not correspond to significant community shifts. We did however identify 3 experimental runs in which the green to red signal ratio from the Chi.Bio passed the threshold over which we predicted a significant shift in community composition, which was also confirmed by plate counting and flow cytometry. The first experimental run was the second replicate of TriComm PVB grown without ampicillin (**Figure 5**, second column); towards the end of the run (from 259 hours until the end of the run), we observed a low GFP to mScarlet signal ratio, indicating that PM was increasing its relative proportion over PL. Consistent with this signal, we observed a 34-fold increase in the ratio of PM over PL colonies with respect to earlier time points, as well as a 3-fold increase in flow cytometry events. The second run for which we observed a shift in community composition was the second replicate of TriComm PN23 grown without colistin (**Figure 5**, third column); at 307 and again at 438 hours the Chi.Bio signal indicated an increased presence of PM over PL, which we confirmed for the first matching plate counting time point with a 25-fold increase at 314 hours. We also observed a significant shift in the other direction, with PL increasing its share of the population over PM, for the second replicate of the TriComm KAN grown in the presence of 0.5 µg/mL of colistin ( **Figure 5**, last column). The GFP to mScarlet ratio measured by the Chi.Bio remained above the threshold for 216 hours between 257 and 473 hours, suggesting that PL increased its share of the population over PM in that time period; this was confirmed by plate counting (with an average of 9-fold increase) and again to a lesser extent by flow cytometry (1.7-fold increase at 356 hours). These results indicate that community composition is only marginally and transiently affected by the selective pressure imposed by antibiotic treatment and its rescue through horizontal gene transfer.

**Figure 5.**
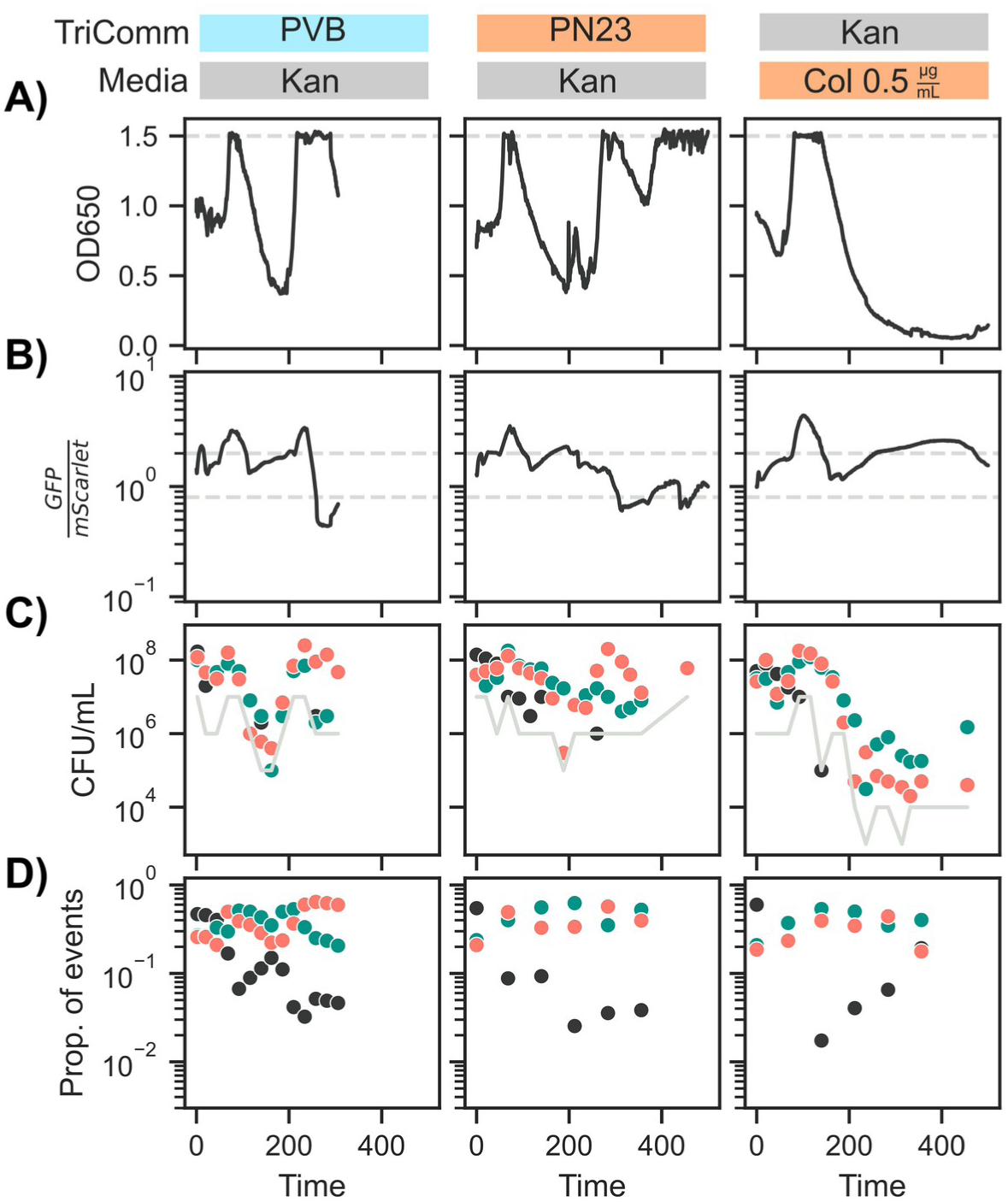
TriComm growth experiments with recorded shifts in the strains’ relative proportions. A) Optical density measurements; the horizontal dashed line indicates the OD level at which the bioreactor switches from chemostat to turbidostat. B) Ratio of green and red fluorescent signals from the bioreactor. Dashed grey horizontal lines indicate the values at which PL (top line) and PM (bottom line) are predicted to be at 1% of the abundance of the other strain. C) Colony forming units for the three TriComm members on non-selective media containing kanamycin. The grey line indicates the limit of detection for the minority strain(s) at each time point. D) Proportion of events belonging to each strain for samples analyzed through flow cytometry.

### Cross-community protection from a non-conjugative pOXA48 plasmid

It has been previously reported that carbapenemases actively degrade carbapenems in a way that provides a protective effect for sensitive strains growing alongside the resistant ones, thus providing a non-genetic path to phenotypic resistance^9–11^. Given that we had included a non-conjugative pOXA48 variant (PVB^23^), we sought to determine whether it provided significant protection to TriComm against ampicillin, which would imply that resistance transfer through conjugation could merely be a selfish side effect of the plasmid backbone, rather than a required property for the community’s survival.

We compared three TriComm variants grown in the presence of 64 µg/mL ampicillin: TriComm KAN, TriComm PVI, and TriComm PVB. In the latter two variants the LM strain harbored the pOXA48 plasmid. As expected, TriComm KAN rapidly collapsed in the presence of ampicillin, as measured by the Chi.Bio optical density sensor and plate counting on non-selective media (*i.e.* without ampicillin, **Figure 6A-B**); conversely, TriComm PVI did not collapse and grew similarly to TriComm KAN grown in the absence of ampicillin ( **Figure 3**). We observed that TriComm PVB’s growth followed a pattern that was intermediate between the other two communities; the community eventually collapsed, but it did so later than TriComm KAN, reaching an OD_650_ of 0.3 after 203 and 245 hours in the two replicates, versus 43 and 34 hours for TriComm KAN (**Figure 6A**). We hypothesized that the ultimate cause of community collapse was a significant reduction in numbers for the PL and PM strains due to the action of ampicillin (**Supplementary Figure 5**), which eventually led LM numbers to decline due to the absence of the lysine and methionine normally provided by PL and PM. Given that the only difference between TriComm KAN and PVB is the presence of the PVB pOXA48 plasmid, we hypothesize that the degradation of ampicillin reduced the effective concentration of the antibiotic in the medium sufficiently to slow down its effect on the sensitive strains. The difference in pOXA48 abundance between the three communities is most evident when using ampicillin selective plating (**Figure 6C**). Read coverage from the sequenced clones confirmed the higher copy number of the PVB variant: the pOXA48 plasmid was present at a median of 7.5 copies per chromosome in the four PVB clones, against a median of 1 copy per chromosome in the six PVI clones (**Supplementary Figure 6**). All four PVB clones carried a single nucleotide change in the *repC* to *repA* intergenic region of pOXA48 (A65388G, 100 bp upstream of *repA*), which was already present in the inoculum LM strain.

**Figure 6.**
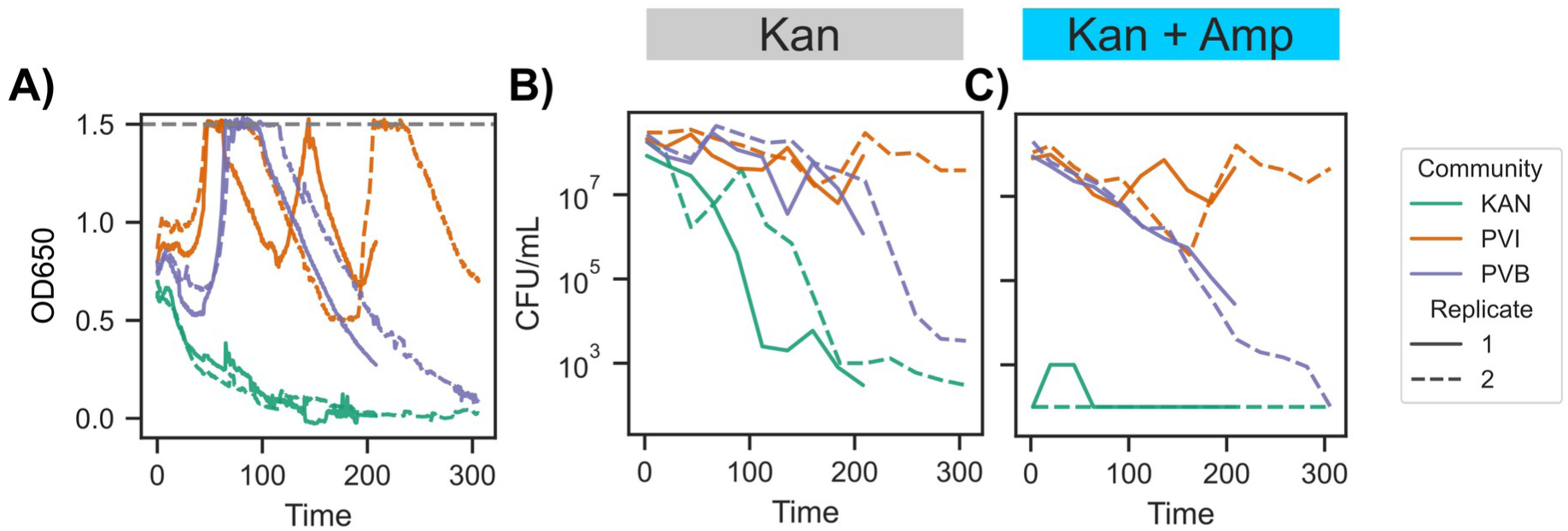
Comparison of TriComm community dynamics during growth in the presence of 64 µg/mL of ampicillin, across three communities and two biological replicates. A) Optical density measurements; the horizontal dashed line indicates the OD level at which the bioreactor switches from chemostat to turbidostat. B) Total colony forming units on non-selective media containing kanamycin. C) Total colony forming units on selective media containing kanamycin+ampicillin.

### A TriComm mathematical model identifies boundaries of AMR plasmid mediated survival under antibiotic treatment

Our empirical measurements of TriComm grown under antibiotic treatment showed that HGT of AMR genes is necessary for community survival. This was especially evident in the case of TriComm PVB, which could only slow down the decline of the community (**Figure 6**). In order to investigate the relationship between the dynamics of HGT and the strength of the antibiotic, and how these two parameters affect a community’s survival, we developed a mathematical model of TriComm (**Figure 7A**, **Methods**). The core of the model included a separate growth rate term (*μ*) for each strain, and a production rate (*α*) for each of the three essential amino acids required for growth. We fit these parameters using TriComm data, specifically plate counting, from growth in the absence of antibiotics; the resulting model captured the oscillations in population size, which were particularly evident from the continuous OD measurements (**Figure 3A**, **Figure 4B** and **Supplementary Figure 4**). The model predicted a gradual dampening of these oscillations, which we confirmed using the longest running TriComm KAN run, not used during parameters fitting (**Supplementary Figure 7**); the model achieved a relatively high correlation with experimental data (Pearson’s *r* 0.33) and with a root mean squared error of 0.50 log₁ CFU/mL and a mean absolute error of 0.41 log₁ CFU/mL.

**Figure 7.**
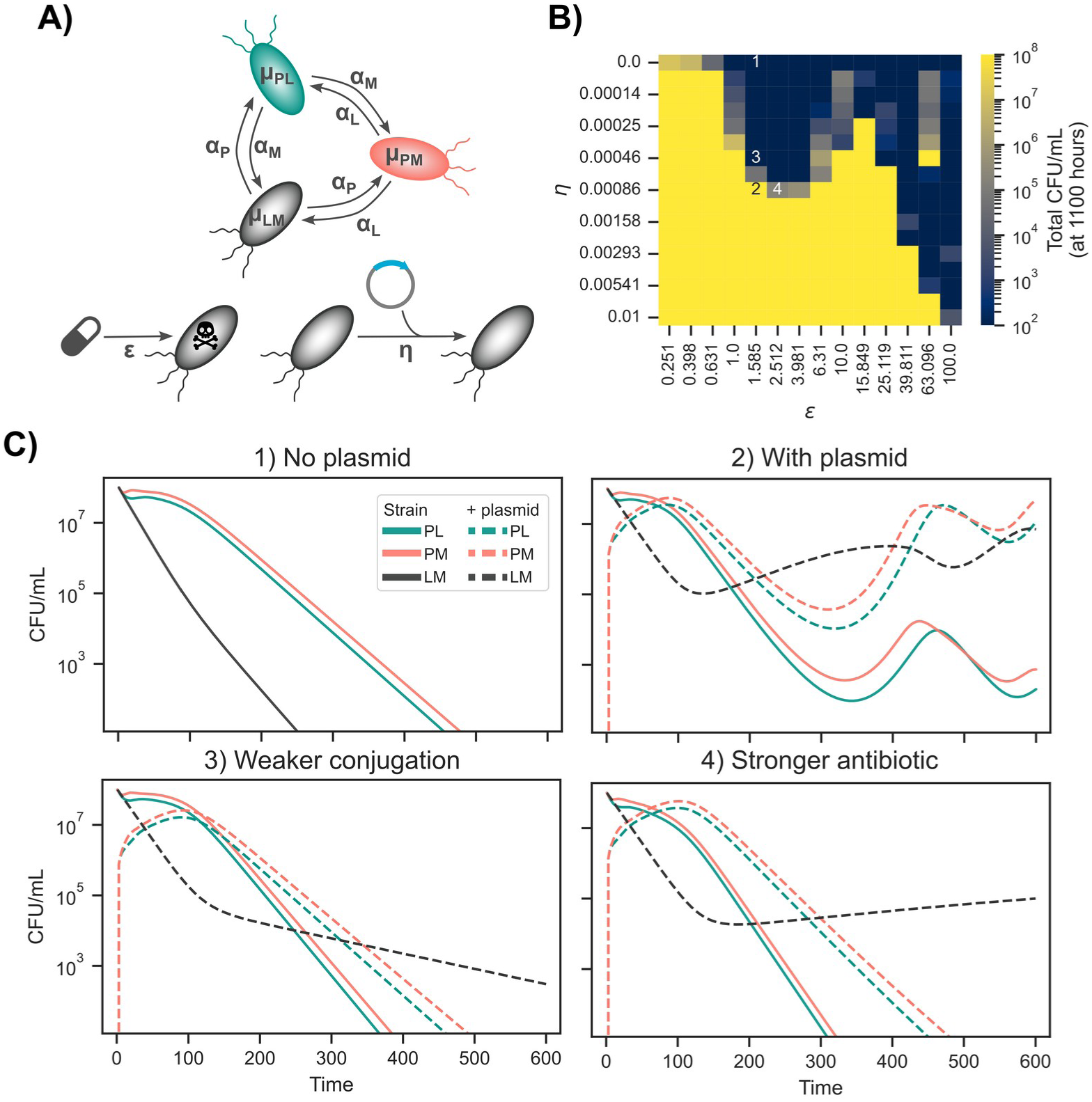
A mathematical model of TriComm and AMR spread via HGT. A) Schematic representation of the model and the main parameters used (see Methods for the full model). B) Two-dimensional parameter sweep on conjugation rate (*η*) and antibiotic potency (*ε*), using the total CFU/mL value at 1,100 hours as a proxy for the community’s survival. C) Model outputs for the four parameter sets highlighted in the previous panel. Solid lines indicate the sensitive strains, dashed lines the same strains harboring the plasmid.

We then extended the model to include antibiotic treatment leading to cell death (*ε*), the effect of which was completely blocked by the presence of the AMR plasmid, and for which its transfer rate between strains was controlled by parameter *η* (**Figure 7A**). All base parameters being equal, the community’s survival in this model is therefore controlled by the combination of these two parameters. We ran a two-dimensional sweep of these two parameters, and used the population size at 1,100 hours as a proxy for community’s survival (**Figure 7B**). We observed an essentially binary outcome in the model’s outputs, with TriComm either surviving or collapsing entirely; the edges in parameter space were particularly sharp, indicating a high sensitivity to changes in parameters. We selected four parameter sets (**Figure 7B**, cells with numbers) and ran the model at those parameter values to inspect the resulting community dynamics (**Figure 7C**). In the absence of the plasmid (set 1), TriComm collapses as expected at almost all the values of *ε* we tested, while a community in which LM has the AMR plasmid at intermediate conjugation rate *η* (set 2) results in a rapid spread of the plasmid to the whole community and thus its eventual survival. Surprisingly, even a small reduction in conjugation rate (set 3) resulting in a slower spread of the plasmid, led to the collapse of the community, as did a stronger effect of the antibiotic (set 4). These results indicate how it may be possible to counteract the spread of AMR plasmids by resorting to more potent drugs, or via the interference with the mechanisms of HGT.

## Discussion

Experimental evolution has been an invaluable tool for understanding bacterial adaptation to antibiotics, but the vast majority of studies have focused on clonal populations in which resistance can only arise via mutation of the existing genetic material^26^. This excludes evolution of resistance via the acquisition of external genetic material (HGT), which is known to be a substantial contributor to the AMR problem. Conjugative plasmids are a special category of HGT that combine the phenotypic impact of their genetic cargo with the ability to self-disseminate within a microbial community. Given the complex dynamics of AMR spread via HGT, we sought to develop an *in vitro* system in which the contribution of each component towards AMR emergence and spread could be measured. Our design objective for this system was to have a stable microbial community, growing in a continuous culture under real-time monitoring.

We used genetic engineering to build our microbial community, thereby removing the effect of background genetic differences between community members from the system. We created our TriComm using *E. coli* BW25113 as a background, and generating three strains, each carrying a separate pair of two gene knockouts in one of three amino acid biosynthetic genes (*metA*, *lysA*, and *pheA*). Previous work had shown how this combination of auxotrophies led to stable growth and interdependency between community members^19^. Through batch culture experiments with defined media, we validated the interdependency of the three TriComm members for growth in minimal media without amino acids. However, we also noticed that a two-member community consisting of the PL and PM strains achieved optical density levels comparable to those of the whole TriComm, although with a longer lag time, a finding we validated in continuous culture. We therefore conclude that the phenylalanine auxotrophy is leaky, which is consistent with previous reports of residual phenylalanine synthesis in *ΔpheA* strains, in which accumulated prephenate converts non-enzymatically to phenylpyruvate at high cell density and mildly acidic pH^27,28^. This partial restoration of the *pheA* deletion may explain why the phenylalanine-producing strain (LM) declines to below 1% of the community during continuous growth experiments, although without going extinct, even after ∼1,000 hours of growth. This suggests that strain LM, while dependent on the other two strains for amino acids, is not essential for the community’s survival; we therefore used it as the initial host for the AMR plasmid.

In order to track community dynamics in real-time, we used the Chi.Bio culturing system, which tracks whole population dynamics via optical density measurements as well as changes in fluorescent signals^22^, providing a way to track the two members (PL and PM) of the community at one-minute intervals during continuous growth. Using calibration experiments with communities with a known proportion of their constitutive members, we validated that the ratio of the two fluorescent signals was able to correctly recapitulate the proportion of GFP-expressing (PL) and mScarlet-expressing (PM) strains, with an estimated limit of detection of 1%. We observed a similar detection limit for the two detection methods we chose to apply to samples taken from the bioreactors, with an exception for both regarding the estimation of the LM strain. Plate counting likely underestimated the abundance of the LM strain given its small colony size, while flow cytometry likely overestimated the LM strain because non-fluorescent PL and PM cells would be counted as belonging to the LM strain. We therefore used the three detection methods together to study TriComm dynamics in subsequent experiments.

We then introduced two conjugative AMR plasmid types into strain LM (pOXA-48 and PN23), and measured community and plasmid dynamics in the presence or absence of antimicrobial treatment. Through selective plating we confirmed that both PVI and PN23 were able to reach all community members via conjugation; we observed transconjugants even in the absence of antibiotics, although at a lower abundance. This indicates that the plasmid fitness cost is not negligible in the absence of a selective advantage for cells harboring it. In the presence of antibiotics, we observed that HGT via conjugation dominated over spontaneous mutations as a mechanism for community survival, with TriComm collapsing in the absence of the plasmid or with a conjugation deficient plasmid (PVB). The only exception was exposure to high colistin concentration (1 µg/mL), in which the PN23 community did not survive. We did not observe any other known AMR mutation in the communities that we subjected to whole genome sequencing, confirming that HGT has the dominant role in spreading AMR.

As previous reports have shown, the *bla*_OXA-48_ gene can provide a public good through periplasmic degradation of its antibiotic substrate^9,11^. This implies that at least in theory conjugation might not be required for community survival. We tested this hypothesis using a non-conjugative natural variant of the PVI plasmid (PVB)^23^. Even though this plasmid had a mutation known to increase its copy number, and thus presumably the rate of degradation of ampicillin, we observed a gradual collapse of the community, although at a reduced pace compared to the plasmid-free KAN community. This difference in the rate of collapse shows how the system we have built is able to provide a quantitative view on community dynamics under treatment and its rescue through HGT.

We developed a simple mathematical model to characterize the ability of HGT to rescue a sensitive obligate community upon antibiotic treatment. To achieve this, we varied both the drug efficacy (*ε*) and the plasmid transfer rate (*η*) and measured the long term (1,100 hours) community levels. The model revealed a sharp boundary between survival and collapse, indicating that altering either parameter alone could determine the community’s outcome. Altering conjugation rates, either chemically through conjugation inhibitors such as unsaturated fatty acids^29^ or relaxase/type-IV-secretion inhibitors^30^, or genetically through deletion or inducible CRISPRi repression of essential transfer genes^31^, would provide a way to test the model’s predictions and develop strategies to block AMR spread via HGT.

As with any experimental system aiming at modeling a complex phenomenon, we are aware of several limitations in our system. TriComm is built from the same genetic background of *E. coli* BW25113, which excludes host range and permissiveness effects in conjugation. The leakiness of the *pheA* deletion reduced the relative importance of the LM strain, leading to an unbalanced community, albeit while still allowing for all three members to coexist over long time periods (**Supplementary Figure 2**). Due to limitations in the available fluorescent markers and their overlapping emission bands, we could only track two of the three members within the Chi.Bio, limiting our ability to track the composition of the full community.

Having established this system as an experimental model to study community dynamics under antimicrobial exposure and its rescue via HGT, we envision using it to further probe this phenomenon and test hypotheses regarding the mechanism of HGT and potential strategies to block it. As suggested by the mathematical model, altering conjugation rates might substantially affect the survival of the community; chemical and genetic tools are available to test this hypothesis in a quantitative manner. As demonstrated with the PVB plasmid, genetic variants within the plasmid, either naturally occurring or introduced through genetic engineering, can be used to map the influence of the transfer machinery and cargo to the effectiveness of community protection and spread of AMR. Expanding TriComm so that a different strain or species is included would also allow studying host barriers and competition in a controlled environment, which would bring this system closer to a more complex environment in which cooperation is not the only interaction. Lastly, the Chi.Bio culturing system allows multiple media inputs and their control, allowing for the study of the effect of changing antibiotic concentration^32,33^. The main advantage of the system would still remain the high level of control over the system parameters, the real time monitoring, and the ability to disentangle the individual factors affecting HGT, which remains a highly relevant factor contributing to the current AMR crisis.

## Methods

### Genetic engineering for the creation of TriComm

Gene knockout mutants of *E. coli* K-12 (BW25113) from the Keio collection^25^ were used in this study. Mutants with independent *lysA*, *metA*, and *pheA* gene disruptions (JW2806-1/KL334, JW3973-1/DL41, and JW2580-1/KA197, respectively) were kindly provided by the Typas lab (EMBL Heidelberg). The knockout alleles came directly from the Keio library and were freshly introduced into the BW25113 background by P1 phage transduction, following the protocol from Prof. Typas’ laboratory^34^.

To generate the second knockout mutation, we used the λ Red recombineering technique as previously described^35,36^, employing the pSIM5 plasmid. A PCR product consisting of a kanamycin resistance (Km) cassette flanked by homology arms corresponding to the regions immediately upstream and downstream of the target gene was generated. The amplified fragment was purified using a PCR clean-up kit and introduced into pSIM5-containing cells via electroporation. Transformants were selected on kanamycin-containing (50 μg/mL) LB agar plates, and successful knockouts were confirmed by colony PCR. The kanamycin cassette was subsequently excised using the FLP recombination system, as described previously.

We used fluorescent plasmids based on the pSEVA backbone (Standard European Vector Architecture)^37^, namely pSEVA2313M (expressing msfGFP) and pSEVA2313S (expressing mScarlet)^38^. Bacterial cells harboring these plasmids were grown overnight in LB medium supplemented with kanamycin (50 µg/mL). Overnight cultures were harvested by centrifugation. Plasmid DNA isolation was performed using the Zyppy plasmid extraction kit (Zymo, Germany). Through electroporation, pSEVA2313M was transformed into strain PL, and pSEVA2313S was transformed into strain PM. Transformant colonies were confirmed by antibiotic selection and whole genome sequencing.

Strains and primers used in this study are listed in **Supplementary Tables 3 and 4**, respectively.

### Phenotypic testing of TriComm members in defined media

Bacterial pre-cultures were streaked from frozen glycerol stock in LB agar with KAN plates, and incubated at 37°C. A single colony was picked from the plate and inoculated into 3 mL liquid LB medium with KAN, shaking at 250 rpm overnight at 37°C. The overnight culture was diluted 1:100 and cultured in 30 mL of fresh LB with KAN. After 4 hours, the cells were harvested and washed twice by centrifugation at 3200 x g in M9 medium (Sigma/Merck, Germany) supplemented with 0.5% glucose, 0.123 mg/mL MgSO_4_, 0.0147 mg/mL CaCl_2_, 0.0028 mg/mL FeSO_4_.7H2O, and 0.01 mg/mL thiamine (vitamin B1). A total of 5 × 10^7^ cells per mL were inoculated into each well containing 200 µL M9 medium with KAN, either without supplements or with a defined amino acid mixture containing 0.01 mg/mL each of L-methionine, L-phenylalanine, and L-lysine monohydrochloride (Sigma/Merck, Germany). Growth was monitored using a Synergy H1 microplate reader over 60 hours, with measurements taken every 20 min, under continuous double-orbital shaking at 237 rpm (4 mm orbit). Experiments were repeated with two independent biological replicates.

### Setup of the Chi.Bio device for the culturing and tracking of TriComm

The Chi.Bio platform (Labmaker, Germany) consists of a control computer, bioreactors and pump boards^22^. The system was set up following the “Operation Manual” V1.2, the “Software Setup Guide” V2.3, and the “Mechanical Assembly Guide” V1.1, which we obtained from the Chi.Bio website (https://chi.bio). Most of the parameters were kept as default setup, but with some exceptions: excitation intensities were set to 0.3 (523 nm LED) and 0.4 (623 nm LED). The pumps were turned on every 2 hours. Pump flow rates were calibrated to values ranging from 0.0043 to 0.005, to get an approximate transfer volume of 600 µL per cycle, thus achieving a dilution factor of 3% per cycle. The efflux pump rate was set to an excess rate of 0.1 to avoid buildup of liquid in the vials and possible spillage. The stirring rate was set to a reduced rate of 0.3 to increase the possibility of prolonged cell-to-cell contact. GFP fluorescence was excited through a 457/35 nm bandpass and detected through a 510/40-nm emission bandpass. mScarlet fluorescence was excited through a 523/70-nm bandpass and detected through a 620/53-nm emission bandpass.

We created a custom Chi.Bio program to run growth experiments both chemostat and turbidostat mode. The chemostat regime was the default setting, actuating a 3% dilution every 2 hours. If the culture’s optical density reached 1.5, the system changed to turbidostat mode, and automatically adjusted the pump rates to maintain the optical density of the culture below the set threshold of 1.5. The custom program and further edits to the default Chi.Bio software are available at: https://github.com/microbial-pangenomes-lab/ChiBio^39^.

### Colony counting

Square LB agar plates were prepared with kanamycin (50 µg/mL), and/or ampicillin (100 µg/mL) or colistin (2 µg/mL), depending on the plasmids carried by the strains being selected for. A volume of 20 µL of Chi.Bio culture was diluted in 180 µL of M9 medium, serially diluted 1:10 to a final dilution of 10^-6^. A volume of 10 µL of each serial dilution was pipetted across 6 tracks per sample, which fitted comfortably onto one half of a square plate. To control the flow of each droplet, the plate was adjusted at an angle between 45° and 90°. Once each track reached the edge of the agar, the plate was placed on the bench and allowed to dry. After drying, plates were incubated at 37°C overnight. Plates were placed inside a blue-light box or left in the fridge for a few days to allow fluorescent protein signal to develop. Colony counts were linked to each TriComm member based on colony color, and CFU/mL was calculated accordingly.

### Flow cytometry

Bacterial culture samples collected from the Chi.Bio continuous culturing experiments were fixed prior to flow cytometry analysis. For each sample, 150 µL of culture was centrifuged at 3200 x g and washed twice with PBS (Roth, Germany), then resuspended in 150 µL of 2% paraformaldehyde (PFA) (Sigma/Merck, Germany) solution and incubated for 1 hour at room temperature in the dark. Samples were subsequently washed and resuspended in PBS, then diluted 100-fold in PBS prior to acquisition.

Samples were run on a Bio-Rad ZE5 Cell Analyzer, with 60 µL acquired per sample at a flow rate of 1 µL/s, yielding between 20,000 and 200,000 cells per sample. To discriminate bacterial cells from background particles, the cytometer was configured with the Small Particle Detector module, using the 405 nm laser for forward scatter and the 488 nm laser for side scatter. GFP fluorescence was excited at 488 nm and detected with a 509/24 nm bandpass filter, while mScarlet fluorescence was excited at 561 nm and detected with a 615/24 nm bandpass filter. These values were chosen to closely mimic those used in the bioreactors. Data were analyzed in FlowJo. Bacterial events were gated on a logarithmic forward-scatter versus side-scatter plot, and fluorescence-based gating was then used to determine the relative proportion of each community member within each sample.

### Calibration of detection methods with defined communities

Bacterial strains were prepared as described in the growth assay section. Based on OD600 and CFU/mL data at 4 hours, all cell concentrations were adjusted to 15×10^7^ cells per mL stock using M9 media. A mixing ratio of 1 thus corresponded to 5×10^7^ cells per mL, while a ratio of 0.01 corresponded to 5×10^5^ cells per mL. We used three methods to assess community composition: fluorescence measurements from the Chi.Bio device, plating on LB agar, and flow cytometry, following the settings described above.

### Continuous culturing of TriComm cultures in the Chi.bio device

Bacterial cultures were prepared as described in the growth assay section. All cell concentrations were adjusted to 5×10⁷ cells/mL using M9 medium without amino acids. Each vial contained a total media volume of 21 mL. The Chi.Bio device was set up as described above to run chemostat and turbidostat modes together. We collected 400 µL of samples daily for storage and analysis by antibiotic selective plating, flow cytometry, and whole-genome sequencing for selected samples.

Chi.Bio raw data files were processed using a Jupyter notebook pipeline. Optical density at 650 nm, and the two fluorescent emission signals were smoothed by taking the median over a sliding 60 minute window. We derived the community composition signal by computing the smoothed GFP emission divided by the smoothed mScarlet emission at each time point.

Colony counts were converted to CFU/mL from the dilution factor and the plated volume, and a per-sample limit of detection was derived from the highest dilution scored. Flow cytometry gates were exported from FlowJo as event counts per strain and converted to proportions of gated events. Rates of decline of plasmid-carrying colonies were estimated by ordinary least squares regression of log₁ CFU/mL on time, excluding time points at the limit of detection. The code used for processing this data is available in a public repository online: https://github.com/microbial-pangenomes-lab/2026_tricomm.

### Whole genome sequencing

Samples for whole genome sequencing were obtained from single colonies picked from selective and non-selective plates and from whole cultures grown overnight from frozen samples (**Supplementary Table 1**). DNA was extracted using the DNeasy Blood & Tissue kit (QIAGEN), and sent for Illumina short read sequencing (Novogene). Reads were analyzed with breseq^40^ v0.39.0 against a composite reference comprising the *E. coli* BW25113 chromosome (RefSeq assembly GCF_000750555.1) and, where relevant, pOXA-48, PN23, pSEVA2313M (msfGFP) and pSEVA2313S (mScarlet). Samples derived from single colonies were analysed in consensus mode, whereas whole-culture samples were analysed in breseq’s polymorphism mode so that segregating mutations could be quantified. The identity of each sequenced isolate was verified from read coverage: the two expected auxotrophy deletions were called from breseq missing-coverage and large-substitution evidence spanning *lysA*, *pheA* and *metA*, and the presence of each plasmid was scored from its mean read depth relative to the mean chromosomal depth, which was also used to estimate plasmid copy number. A variant already present at a frequency of at least 0.5 in the corresponding ancestral clone was treated as pre-existing rather than specific to the analyzed sample. Variant calls falling within the cryptic prophage regions DLP12, e14, Rac, Qin, and CPS-53, as well as the *rrn* operon were excluded to remove low-confidence variant calls. All variant calls across samples are available in **Supplementary Table 5**. Analysis scripts are deposited in a public repository: https://github.com/microbial-pangenomes-lab/2026_tricomm_sequencing.

### Mathematical modeling

The mathematical model was constructed based on the chemical reactions describing the community dynamics (eq. 1). Strain LM grows at a rate of *μ_LM_*, only when lysine (*aa_L_*) and methionine (*aa_M_*) are available in the growth medium, and produces phenylalanine (*aa_P_*) at a rate of α_LM_. Strains PM and PL follow analogous dynamics, while population loss for all strains is driven by the dilution rate δ.

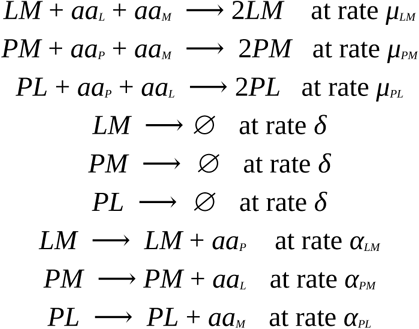

The effect of the antibiotic is described by the parameter ε. The capacity of a resistant strain to transfer the plasmid conferring antibiotic resistance is described by η, which depends on the total population of the three resistant strains.

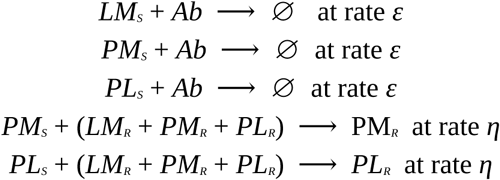

Antibiotic concentration decreases due to dilution δ and natural degradation γ (Eq. 3), while being constant supplied at a constant rate of δ*Ab_init_*.

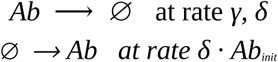

System dynamics are modeled using ten differential equations within a generalized Lotka–Volterra framework based on the chemical reactions in equations 1-4, tracking time-dependent changes in strain populations, amino acid concentrations and antibiotic strength (Eq. 5). According to the model structure, each strain population changes according to the availability of the amino acid concentration and the dilution steps, as well as the antibiotic concentration, its impact on sensitive strains, and the presence of the antibiotic-resistance plasmid. On the other hand, each amino acid concentration depends on its production by the corresponding strain and its utilization by the strains that rely on it. Subscripts *S* and *R* denote sensitive and resistant strains, respectively, with total strain population expressed as the sum of both subpopulations, *LM_R+S_*, *PM_R+S_* and *PL_R+S_*.

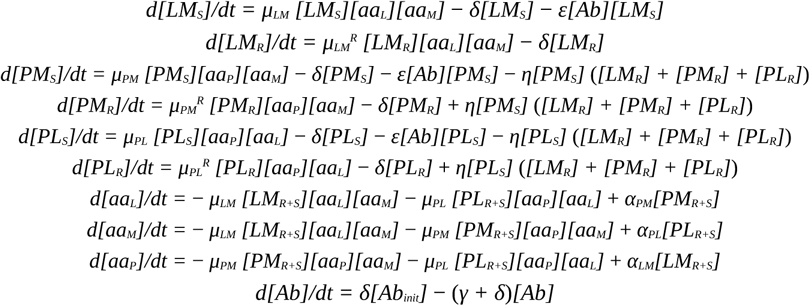

Model parameters were estimated by fitting the equations to experimental data, using the mean CFU/mL trajectories of the three strains from the KAN TriComm vials (replicate 2, KAN plate counts), averaged across the four vials and linearly interpolated onto a denser time grid. This estimation was carried out in MATLAB R2025a using the SIMPLEXL optimization algorithm. The objective function was defined as the sum of the mean squared errors associated to each strain population. Due to the limited number of experimental data points, the time series were interpolated to improve the accuracy of the error estimation, using the MATLAB built-in *interp1* function. In total, seven parameters were fitted: the growth rate of the three strains (μ_LM_, μ_PL_, μ_PM_) the production rates of each amino acid (aa_L_, aa_M_, aa_P_) and the dilution rate (δ). Confidence intervals of 95% for each parameter were calculated using the MATLAB built-in *nlparci* function (**Supplementary Table 6**).

To evaluate community survival under varying antibiotic strength and resistance transfer capabilities, we conducted a two-dimensional parameter sweep across the antibiotic killing rate (ε) and the plasmid conjugation rate (η). Parameter ranges were set to ε ∈ [0.251, 100.0] h^-1^ and η ∈ [1×10⁻⁴, 1×10⁻²] h^-1^, log-spaced across 16 values, together with a η = 0 no-conjugation control (14 × 17 combinations in total). Each simulation was initialized with the experimental initial strain abundances and integrated numerically up to t=1100 h (dt = 4 h) using GNU Octave’s lsode solver. Total community survival was quantified by recording the final combined strain population (expressed in CFU/mL) at t=1100 h. Four specific parameter combinations were further selected from this parameter space to analyze individual strain time-course trajectories.

## Data availability

All data and related analysis scripts are deposited in two public repositories (https://github.com/microbial-pangenomes-lab/2026_tricomm and https://github.com/microbial-pangenomes-lab/2026_tricomm_sequencing), with specific releases archived in Zenodo^41,42^. Raw sequencing reads have been deposited in the European Nucleotide Archive (ENA) under accession number PRJEB127418.

## Author contributions

M.G. developed the research concept together with T.H.V. T.H.V. ran all laboratory experiments, with help from L.L. for genetic engineering, C.A. for flow cytometry, A.M. for colony counting, and K.G. for operating Chi.Bio experiments. T.A.V., E.S., F.D.P., and M.F. developed the mathematical model. M.G. analyzed the experimental and sequencing data with help from H.T.V.. M.G. and T.H.V. wrote the manuscript. All authors revised the initial draft and approved the final version.

## Acknowledgments

We thank Alvaro San Millan for gifting the PVI and PVB plasmids and helpful discussions. We thank Craig MacLean for gifting the PN23 plasmids. We thank Till Strowig for the helpful discussions. We thank Alison Mather and Mark Webber (Quadram Institute) for suggesting the name TriComm and helpful discussions. T.H.V., A.M., K.G., and M.G. were supported by the Deutsche Forschungsgemeinschaft (DFG, German Research Foundation) under Germany’s Excellence Strategy – EXC 2155 (Project No. 390874280). T.H.V. and K.G. were further supported by the Hannover Biomedical Research School (HBRS), and the Centre for Infection Biology (ZIB). T.H.V. was also supported by the Graduate School Scholarship Programme (GSSP) from the German Academic Exchange Service (DAAD). K.G. was funded by zukunft.niedersachsen, the joint science funding program of the Lower Saxony Ministry of Science and Culture and the Volkswagen Foundation (BaKlimON; reference ZN4791). M.F. and T.A.V. were supported by two PRIN grants from the Italian Minister of University and Research, *i.e.* Microbiome Assembly: A multilevel approach to predict establishment and variation of microbiomes (Grant n. 2022Z88RK4) and EXPLORE: EXploiting pathogens PLOidy to fight drug REsistance (Grant n. P2022AB5TY).

