## Supplementary Information for "Development of a stable microbial community for the study of plasmid-mediated antimicrobial resistance in a continuous culturing device"

### Supplementary Materials

### Supplementary Figures

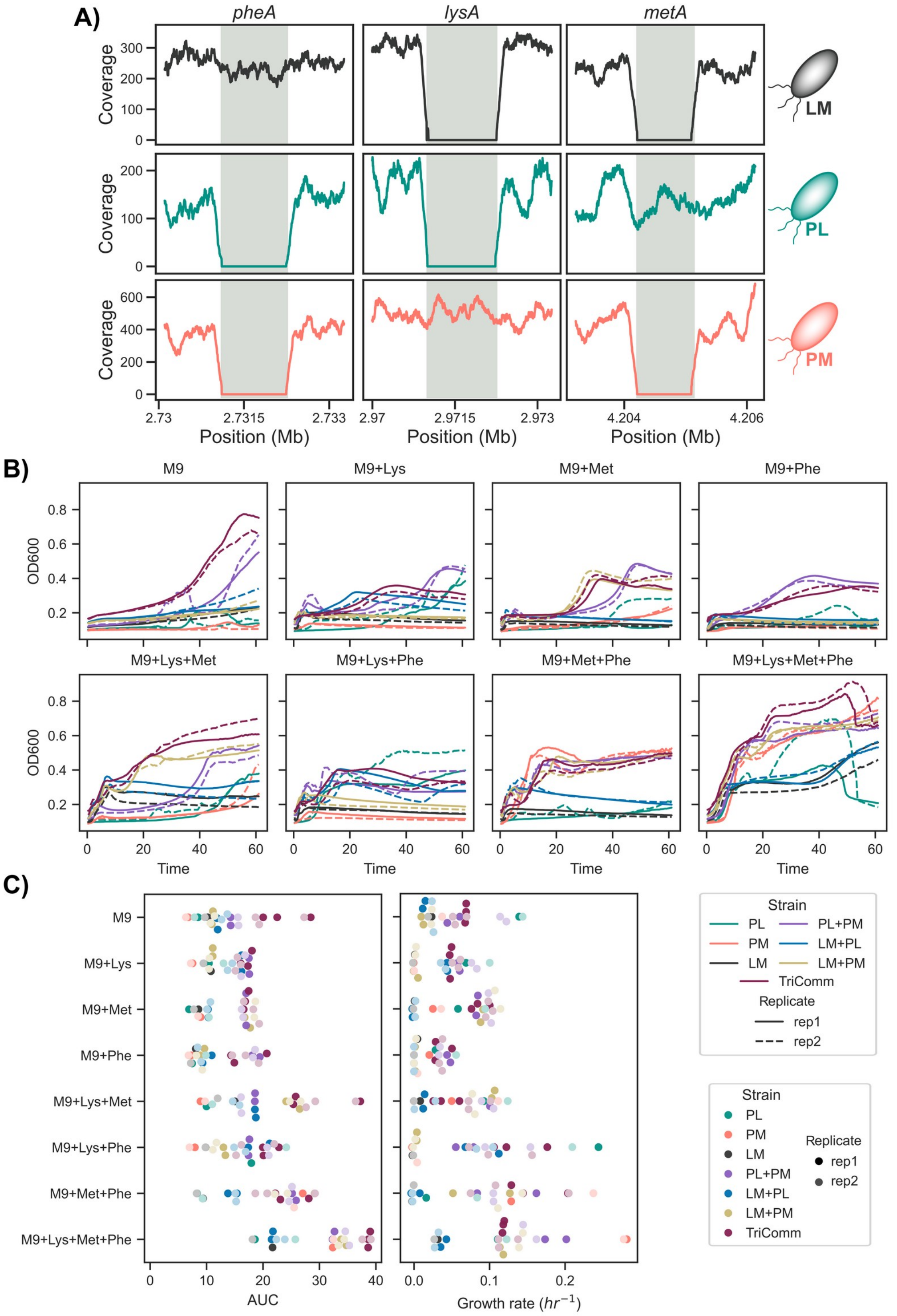


**Supplementary Figure 1. Genetic and phenotypic characterization of TriComm.** A) Read coverage after mapping to the *E. coli* BW25113 reference in the regions corresponding to the three target aminoacid biosynthetic genes in the three TriComm strains. The grey boxes indicate the location of each target gene. B) Growth curves for each strain and community across each specific media formulation. Average growth curves across technical replicates for the two biological replicates. Confidence intervals are omitted for readability. C) Area under the curve (AUC) and growth rates for each strain and community grown across media compositions.

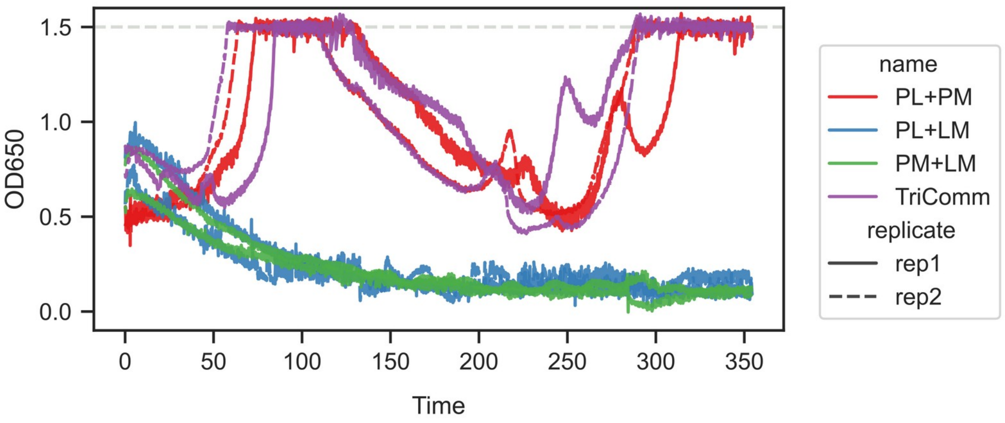


**Supplementary Figure 2. Growth of 2-member communities and TriComm in continuous culture.** Two biological replicates are shown for each tested community.

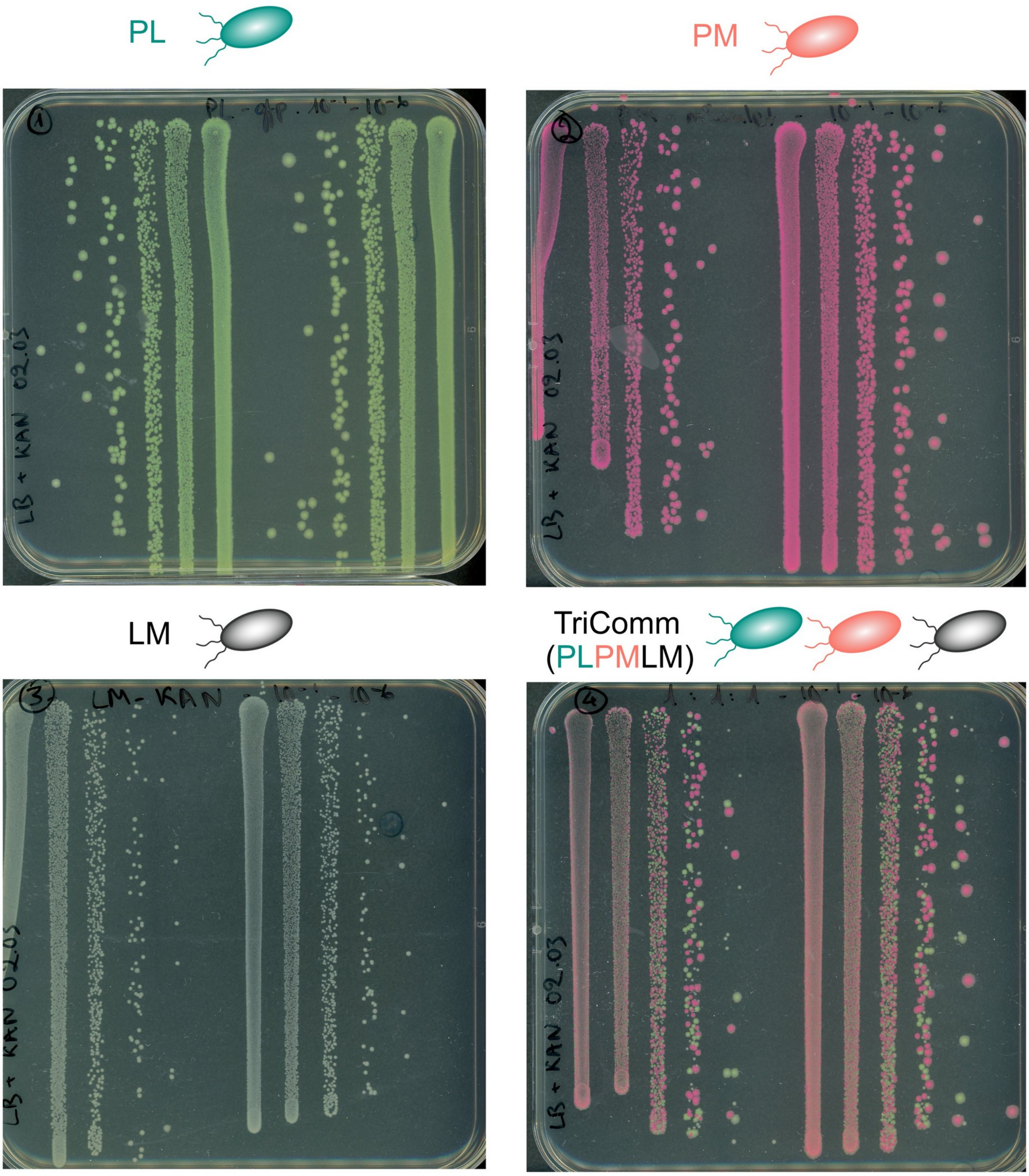


**Supplementary Figure 3. Strain LM develops smaller colonies than strains PM and PL.** Square Petri plates with sequential dilutions from the experiments described in **Figure 2**.

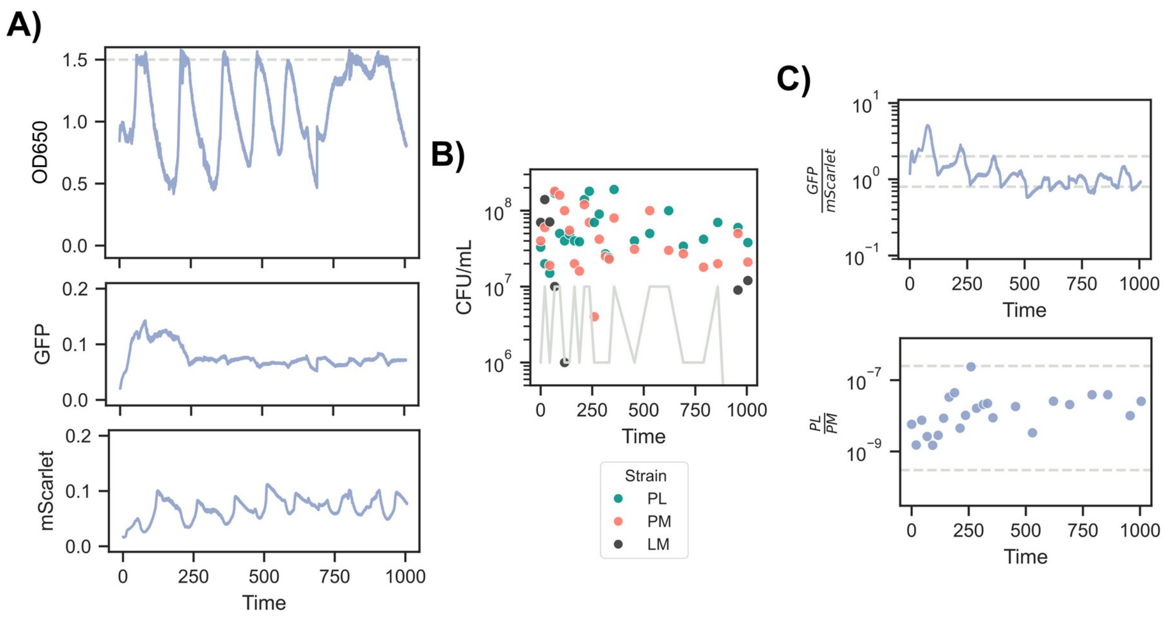


**Supplementary Figure 4. TriComm community dynamics during long continuous growth.** A) Optical density and fluorescence measurements taken at one minute intervals by the Chi.Bio device. The horizontal dashed line indicates the OD level at which the bioreactor switches from chemostat to turbidostat. B) Colony forming units for the three strains from daily samples; the grey line indicates the limit of detection for the minority strain. C) Ratio of green and red signals from the bioreactor (top) and plate counting (bottom). Dashed grey horizontal lines indicate the values at which PL (top line) and PM (bottom line) are predicted to be at 1% of the abundance of the other strain.

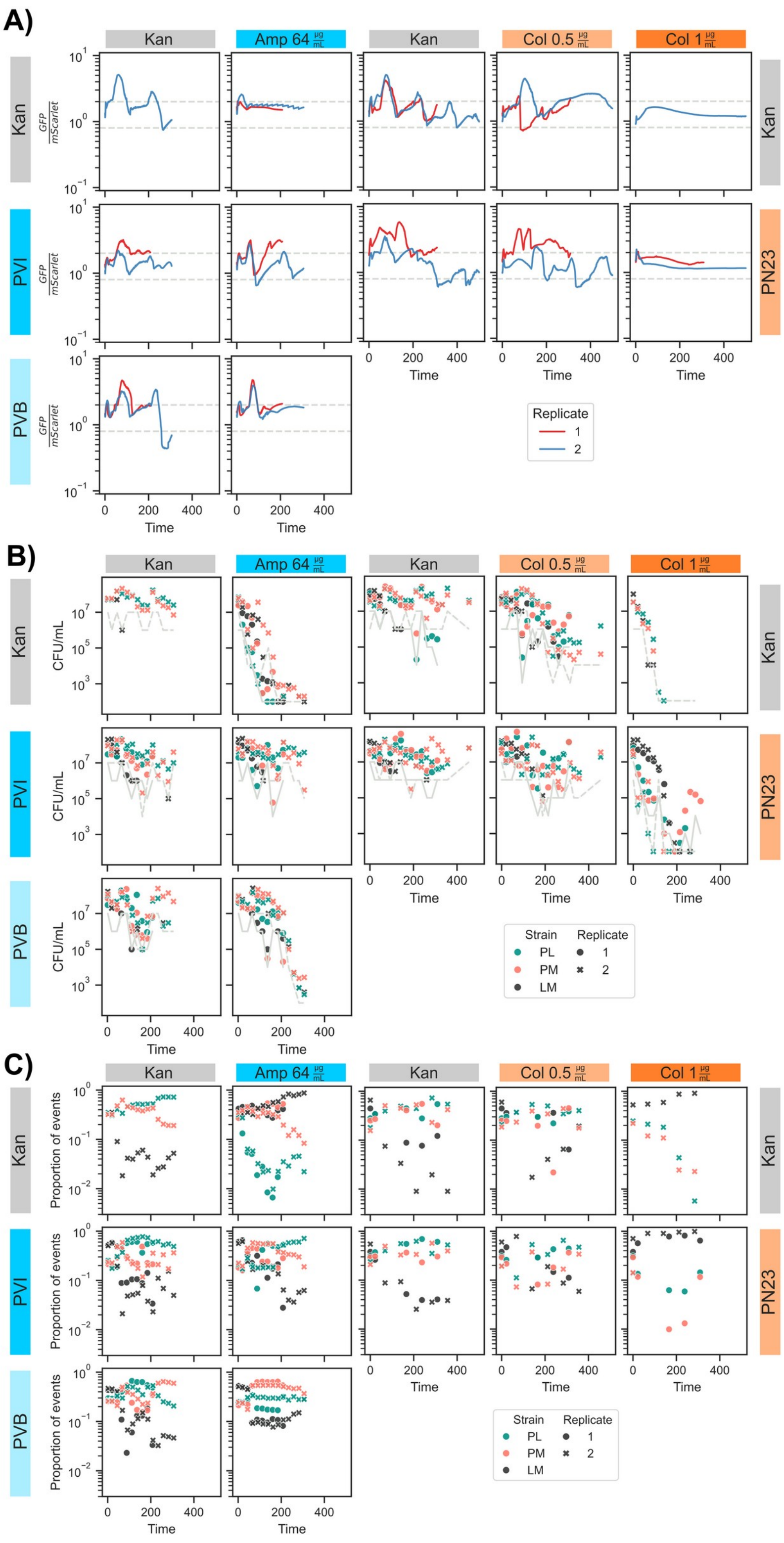


**Supplementary Figure 5. TriComm community dynamics under different selective pressures and conjugation regimes.** A) Ratio of green and red fluorescent signals from the bioreactor. Dashed grey horizontal lines indicate the values at which PL (top line) and PM (bottom line) are predicted to be at 1% of the abundance of the other strain. B) Colony forming units for the three TriComm members on non-selective media containing kanamycin.

The grey line indicates the limit of detection for the minority strain(s) at each time point. C) Proportion of events belonging to each strain for samples analyzed through flow cytometry.

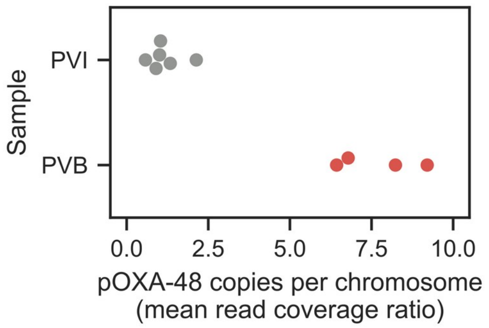


**Supplementary Figure 6. Ratio of reads mapped to the pOXA48 plasmid over those mapped to the chromosome from PVI and PVB clonal samples.**

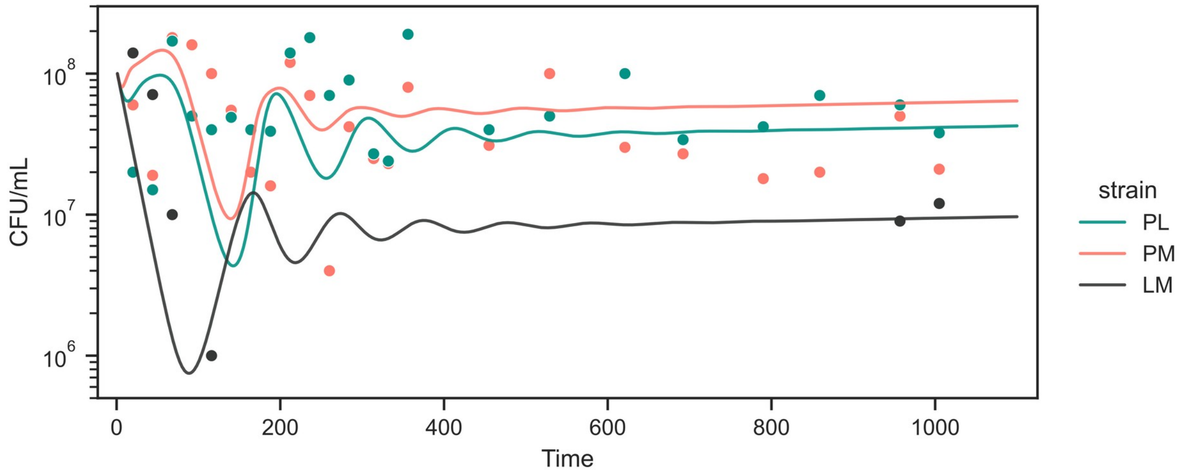


**Supplementary Figure 7. Comparison of the mathematical model (solid lines) with the plate counting data (dots) for a TriComm KAN community grown in the absence of antibiotics.**

### Supplementary Discussion

#### Adaptive mutations unrelated to antimicrobial resistance

We did not identify genetic variants that were either clearly associated with resistance to ampicillin or colistin, nor potentially associated with resistance and found exclusively on experimental runs treated with the two antibiotics. We therefore investigated the potential relation of the remaining de-novo mutations to the aminoacid auxotrophies or to growth in liquid cultures.

##### Adaptaptation to auxotrophic community growth

Several genetic variants can be linked to ameliorating TriComm's auxotrophies. The strongest candidate is a truncating variant in *pykF* (K13), fixed in a PL clone (pOXA48_rep2/M5_PL_clone2, TriComm PVI 16xMIC AMP). PykF consumes phosphoenolpyruvate, which is also a precursor for the shikimate pathway, leading to chorismate and aromatic amino acids. Loss of *pykF* can therefore redirect carbon flux toward aromatic biosynthesis, and thus to availability of phenylalanine to the *pheA* mutants[1].

Among other candidate variants is the G2242020C substitution in the *lysP–yieE* intergenic region, which increased from 0.18 to 0.56 during the PN23 experiment and was fixed in a PL clone (TriComm PN23 1xMIC COL, N=3: PN23_rep2/M3_314h, PN23_rep2/M3_455h, and PN23_rep2/M3_PL_clone1). This variant may affect the uptake of lysine via LysP for the PL and LM strains. A separate +T insertion in the same region occurred in sample pOXA48_rep1/M5_198h (TriComm PVI 16xMIC AMP). Concerning lysine, we found two distinct variants in the *lysO–aqpZ* intergenic region: A910669G (pOXA48_rep2/M4_306h, TriComm PVI no drug) and C910664T (pOXA48_rep2/M5_306h, TriComm PVI 16xMIC AMP). Since LysO is a lysine exporter, these variants could affect lysine release to the community.

**Adaptation to sustained growth in liquid culture**

We flagged several genetic variants for their potential role in favoring adaptation to growth in liquid media, although with varying levels of confidence. Several candidates were involved in respiration or nutrient acquisition, including *cyoA* N172S (pOXA48_rep1/M7_198h, TriComm PVB 16xMIC AMP), *nfuA* H126R (pOXA48_rep1/M5_198h, TriComm PVI 16xMIC AMP), and *pstA* V168A (pOXA48_rep1/M7_198h, TriComm PVB 16xMIC AMP). We also identified a large variety of variants affecting envelope-related genes; even though some of these are known to be involved in AMR, we identified them in both treated and untreated samples. The *ompR* substitutions included G148C (N=2: PN23_rep2/M3_314h, TriComm PN23 1xMIC COL; pOXA48_rep1/M5_198h, TriComm PVI 16xMIC AMP), A167E (PN23_rep2/M3_455h, TriComm PN23 1xMIC COL), T149K (N=3: PN23_rep2/M3_455h, TriComm PN23 1xMIC COL; PN23_rep2/M3_PM_clone1, TriComm PN23 1xMIC COL; pOXA48_rep1/M5_198h, TriComm PVI 16xMIC AMP), K170N (PN23_rep1/M6_PM_clone1, TriComm PN23 1xMIC COL) and A167T (pOXA48_rep2/M4_306h, TriComm PVI no drug). Variants were also detected near *ompF*, including C982566A (PN23_rep2/M3_314h, TriComm PN23 1xMIC COL) and T982560A (N=3: PN23_rep2/M3_PL_clone1, TriComm PN23 1xMIC COL; pOXA48_rep1/M6_198h, TriComm PVB no drug; pOXA48_rep2/M6_306h, TriComm PVB no drug), as well as near *ompC* (pOXA48_rep2/M0_PL_clone1, TriComm KAN no drug). The *cpxA* A112D substitution occurred in two LM clones (N=2: pOXA48_rep1/M4_LM_clone1, TriComm PVI no drug; pOXA48_rep1/M5_LM_M9, TriComm PVI 16xMIC AMP). The role of these genes in evelope stress and porin expression makes them potential candidates for long term adaptation to continuous liquid culture[2,3].


1. Meza, E., Becker, J., Bolivar, F., Gosset, G. & Wittmann, C. Consequences of phosphoenolpyruvate:sugar phosphotranferase system and pyruvate kinase isozymes inactivation in central carbon metabolism flux distribution in Escherichia coli. *Microb. Cell Factories* **11**, 127 (2012).
2. Zhang, E. & Ferenci, T. OmpF changes and the complexity of Escherichia coli adaptation to prolonged lactose limitation. *FEMS Microbiol. Lett.* **176**, 395–401 (1999).
3. Liu, X. & Ferenci, T. Regulation of Porin-Mediated Outer Membrane Permeability by Nutrient Limitation in Escherichia coli. *J. Bacteriol.* **180**, 3917–3922 (1998).
